# eccDNA Atlas: a comprehensive resource of eccDNA catalog

**DOI:** 10.1101/2022.11.06.515328

**Authors:** Tengwei Zhong, Wenqing Wang, Houyan Liu, Maolin Zeng, Xinyu Zhao, Zhiyun Guo

## Abstract

Extrachromosomal circular DNA (eccDNA) represents a large category of nonmitochondrial and nonplasmid circular extrachromosomal DNA playing an indispensable role in various aspects such as tumorigenesis, immune responses, liquid biopsy, etc. However, characteristic and functions of eccDNA are fragmented, hiding behind abundant literatures and massive whole-genome sequencing (WGS) data, which have not been sufficiently used for identification of eccDNAs. Therefore, establishing an integrated repository portal is essential for identifying and analyzing eccDNAs. Here, we developed eccDNA Atlas (http://lcbb.swjtu.edu.cn/eccDNAatlas), a user-friendly database of eccDNAs that aims to provide a high-quality and integrated resource for browsing, searching and analyzing eccDNAs from multiple species. eccDNA Atlas currently containing 630,434 eccDNAs and 7,774 ecDNAs manually curated from literatures and 1,105 ecDNAs predicted by AmpliconArchitect based on WGS data involved in 66 diseases, 57 tissue and 319 cell lines. The content of each eccDNA entry included multiple aspects such as sequence, disease, function, characteristic, validation strategies, etc. Furthermore, abundant annotations and analyzing utilities were provided to explore existed eccDNAs in eccDNA Atlas or user-defined eccDNAs including oncogenes, typical enhancers, super enhancers, gene expression, survival, and genome visualization. Based on these resources, eccDNA Atlas will significantly improve our understanding of eccDNAs and serve as an important catalyst for future research of eccDNAs.

## Introduction

Extrachromosomal circular DNA (eccDNA) is a nonmitochondrial and nonplasmid circular structured DNA which is widely found in eukaryotes[1]. Up to now, eccDNAs with different nomenclatures have been discovered in a variety of species such as ecDNA (previous refers to double minute)[2], small polydispersed circular DNA (spcDNA)[3-6], microDNA[7], telomeric circles or t-circles[8, 9], extrachromosomal ribosomal circles (ERC)[10, 11], etc. Among them, ecDNA has been considered as the most crucial type of eccDNA for wide study. EcDNA (extrachromosomal DNA) refers to the long extrachromosomal circular DNA (168Kb∼5Mb, with an average of 1.26Mb) which mainly exists in tumor and can be observed in the metaphase of mitosis[12, 13]. EcDNA appears as monomers or clusters[14], and usually carries a variety of oncogenes such as DHFR, EGFR, MYC[15]. EcDNA can cause high oncogene amplification by random division or by enhancer regulation, promoting drug resistance, tumor heterogeneity and even driving poor outcome for patients[2, 16-18]. In addition, the other types of eccDNA except ecDNA (hereinafter referred to as eccDNA) have also been confirmed to have crucial effect s in eukaryotes. Unlike ecDNA, which is only present in tumors, eccDNA can be widely founded in different cell types including health and other disease cells[19]. Although eccDNA is generally smaller than ecDNA, studies have shown that they also sometimes carry and express genes or miRNA to promote tumorigenesis[20]. Previous studies showed that eccDNA could be released into circulation such as plasma or serum, suggesting that they may serve as a biomarker for disease[21]. Furthermore, eccDNAs with smaller size can function as potent innate immunostimulants in a manner that is independent of eccDNA sequence but dependent on eccDNA circularity and the cytosolic DNA sensor sting[22].

How to accurately identify and characterize eccDNAs/ecDNAs is the key point to study their functions. Previously, diverse methods have been developed to perform relevant researches including sedimentation in a sucrose gradient[19], electron microscopy[19], light microscopy[4], southern blotting[23], probe hybridization, florescence *in situ* hybridization (FISH)[24] and 2D gel electrophoresis[25]. In recent years next generation sequencing (NGS) technologies have been developed, especially the improvement of whole-genome sequencing technology lead to the emergence of new methods for identification of eccDNAs/ecDNAs. For example, AmpliconArchitect currently has been considered as the most robust and feasible tool for predicting ecDNA focally amplified regions based on WGS data[26]. Moreover, another widely used method called Circle-Seq has been developed for isolation and sequencing of eccDNA on a genomic scale[27]. The emergence of these methods mentioned above has generated vast amounts of eccDNA data and greatly promoted the research of eccDNA.

Nowadays, lots of wet and dry experiments have been performed to identify and clarify the functions of eccDNAs/ecDNAs across multiple species, especially after the completion of the TCGA and pan-genome project[28]. Characteristic and functions of eccDNAs/ecDNAs are fragmented and hide behind almost countless literatures. Besides, a large number of WGS data was released by numerous studies. Therefore, integrating the above resources to build a database to store eccDNAs/ecDNAs data is very important for the follow-up study of eccDNAs/ecDNAs. To date, two excellent eccDNA databases have been developed to explore this information. CircleBase[29] extracted eccDNAs from 13 literatures and perform six annotations based on the genomic and epigenetics data of eccDNAs. eccDNAdb[30] identified human eccDNAs by AmpliconArchitect from WGS data (131 samples from Turner *et al*. study and 60 tumor samples from SRA). However, there two databases are not comprehensive enough, as they only included eccDNAs from human species, and the WGS data and literatures used are insufficient. Therefore, it is necessary to integrate as much as possible existing eccDNA resources in literatures and predict new eccDNAs based on available WGS data for the subsequent research of eccDNA.

In this study, we developed a database, called eccDNA Atlas, to extend resources of eccDNAs/ecDNAs. We extracted eccDNAs/ecDNAs from 3,636 literatures using eccDNAs/ecDNAs keywords and finally obtained 638,208 eccDNAs (7,774 ecDNAs) from seven species. Furthermore, we downloaded WGS data of 319 tumor samples for identifying ecDNAs by AmpliconArchitect, if the literature already has identified eccDNAs/ecDNAs which predicted based on WGS, we will collect these eccDNAs/ecDNAs according to literature form (for example, 131 samples with WGS data from Turner *et al*. study used by eccDNAdb database). Finally, 1,105 ecDNAs were identified. Besides, we obtained 1,325 oncogenes, 116,660 typical enhancers and 309,844 super enhancers on these eccDNAs/ecDNAs and provided utilities for analysis and genome visualization. In addition, a user-friendly web interface was built and split into eight main pages: (I) Browse, (II) Search, (III) Analysis, (IV) Genome-browser, (V) Download, (VI) Statistics, (VII) Submit and (VIII) Help.

## Materials and methods

### Data collection

#### Data collection from literature

Due to the confusion of eccDNA nomenclatures, the eccDNA names appeared in literatures are very ambiguous. Therefore, we manually reviewed all eccDNA aliases and types names (full name or abbreviation) of eccDNA in literatures and class them into these categories as follows: ecDNA, extrachromosomal DNA, eccDNA, extrachromosomal circular DNA, microDNA, spcDNA, extrachromosomal ribosomal circles and telomeric circles/t-circles. To ensure the highest quality in data collection process, all eccDNA entries were manually collected by the following steps. 1) a list of keywords with restrictions were searched in the PubMed. For example, the search keywords for ecDNA was ‘ (“extrachromosomal DNA” * [Title/Abstract], “ecDNA”, “extrachromosomal DNA”) ‘, and the search keywords for eccDNA was ‘ (“eccDNA” * [Title/Abstract], “eccDNA”, “extrachromosomal circular DNA”, “microDNA”, “spcDNA”, “t-circles”, “telomeric circles”, “extrachromosomal ribosomal circles”) ‘. Therefore, 7,212 literatures were obtained through the above search strategy. 2) After manually removing the duplicates and review literatures, 3,636 literatures as the final literatures were extracted for further eccDNA collection. 3) By manually curating from 3,636 literatures, we found that there are a large number of extrachromosomal DNAs was classified into plasmid DNA/telomeres. Finally, 814 literatures belonging to the eccDNA research were selected for eccDNA data extraction according to the following information catalog: (I) Species and eccDNA type; (II) Chromosomal localization of eccDNAs/ecDNAs; (III) Sample condition and disease; (IV) Validation strategies and sequencing library types; (V) The function/characteristic of eccDNAs/ecDNAs; (VI) PMID and publish date and (VII) Other information. All information was double checked and the names of cell lines were also checked according to nomenclature of cell lines.

#### Prediction of ecDNA based on WGS data

In order to obtain more comprehensive eccDNAs/ecDNAs data, available WGS data from 319 tumor samples which has not been used to predict eccDNAs/ecDNAs in previous studies (eg. WGS data from Turner et al. study) were downloaded from SRA database. AmpliconArchitect was employed for identification of eccDNAs/ecDNAs amplicons according following criteria: genomic segments >10 kb with copy numbers (CNs) >4. Finally, the predicted amplicons were classified using AA classifier. Figure 1 shows the flowchart of the database construction process.

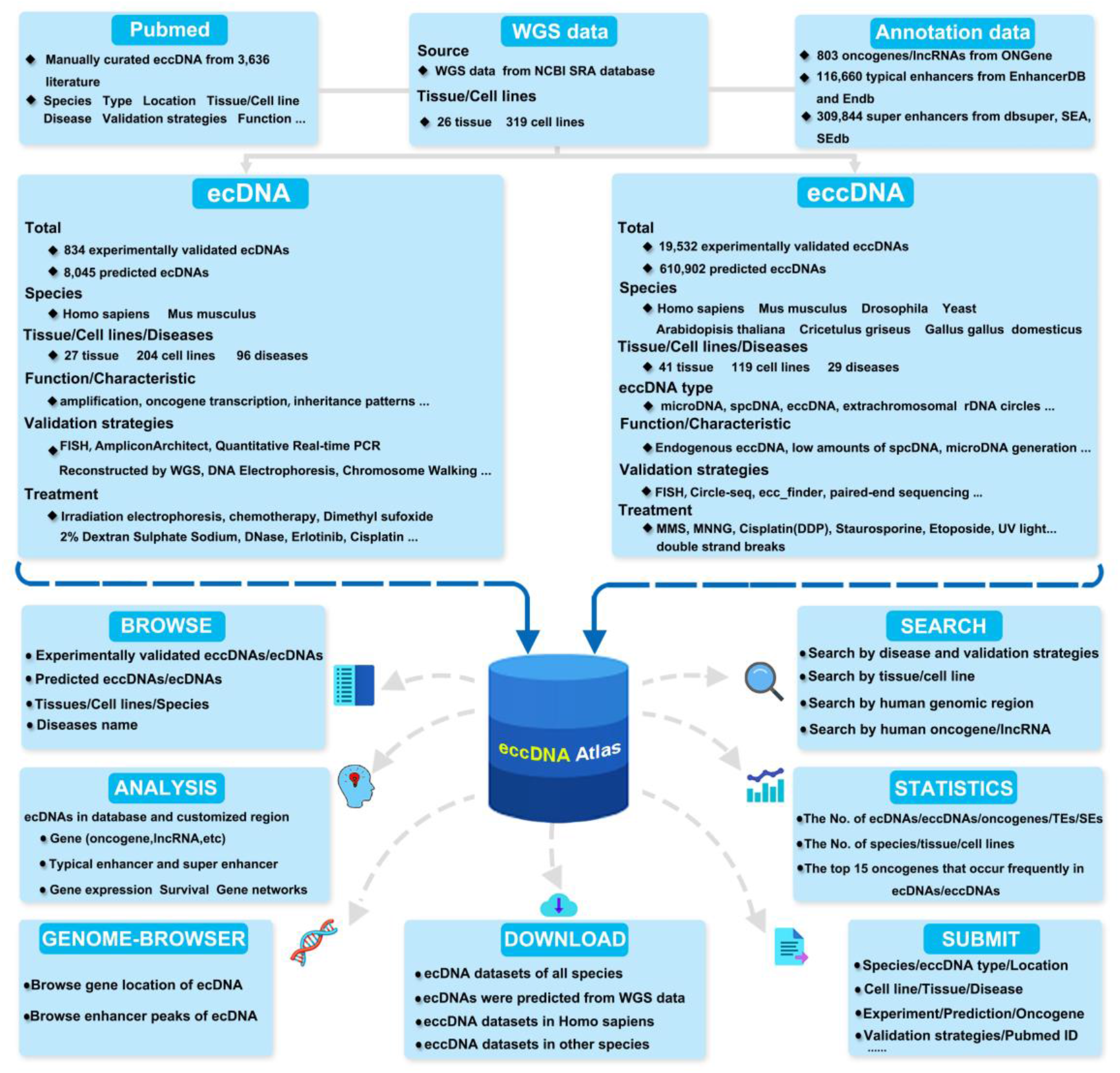

### ecDNA/eccDNA annotation and visualization

The R package (biomartR) was used to gather genomic information for genome version hg19, and a total of 803 oncogenes were download from ONGene[31] (http://www.ongene.bioinfo-minzhao.org/index.html). A total of 116,660 typical enhancers were downloaded from ENdb[32] and EnhancerDB[33] database, and 309,844 super enhancers were downloaded from dbsuper[34], SEA[35], and SEdb[36] database. If a regulatory element overlaps with an eccDNA/ecDNA region, it was annotated as an eccDNA/ecDNA regulatory element. For the convenience of users, in addition to providing the ecDNA annotations already in the database, we also allow users to input customized chromosome positions for annotation. UCSC browser was used to provide visualization annotation of current eccDNA/ecDNA region and user-defined eccDNA/ecDNA region, such as histone modification signals, mutations, sequence conservatism, etc. In order to provide more gene information, the GEPIA[37], GeneCard[38], and String[39] databases of external links were provided for gene expression, function, survival and regulatory network exploring, etc.

### System design and implementation

The eccDNA Atlas website runs on a Nginx Web server (https://nginx.org/). The database was developed using MySQL 5.7.27 (http://www.mysql.com). PHP7.2.30 (http://www.php.net) was used for server-side scripting. The eccDNA Atlas web interface was built using Bootstrap v3.3.7 (https://v3.bootcss.com) and JQuery v2.1.1 (http://jquery.com). ECharts(http://echarts.baidu.com) was used as a graphical visualization framework. We recommend using the latest versions of Firefox and Google Chrome for the best experience.

## Results

### Database statistics

Currently, two types of data were involved in eccDNA Atlas: (I) 630,434 eccDNAs and 7,774 ecDNAs manually curated from literatures; (II) 1,105 ecDNAs identified by AmpliconArchitect based on public WGS data. The detailed statistics were listed in the **Table 1**.

**Table 1.** Statistics of eccDNAs and ecDNAs in eccDNA Atlas.

| Type | eccDNA | ecDNA |
| --- | --- | --- |
| Curated from literatures | 630,434 | 7,774 |
| Identified by WGS | 414,795 | 8,293 |
| Species | 7 | 2 |
| Normal sample | 191,138 | - |
| Disease sample | 439,296 | 8,879 |
| Cell line | 571,107 (from 119 cell lines) | 1,951 (from 204 cell lines) |
| Tissue | 59,327 (from 41 tissue) | 6,928 (from 27 tissue) |
| Disease | 29 | 96 |
| Experimentally validated | 19,532 | 834 |
| Prediction | 610,902 | 8,045 |
| Oncogene | 680 | 645 |
| Typical enhancer | 116,660 |  |
| Super enhancer | 309,844 |  |

### Web interface and usage

We developed a user-friendly web interface to help users to browse, search, analysis, download and submit the eccDNA/ecDNA data. The web interface was split into the following eight main pages: (I) Browse, (II) Search, (III) Analysis, (IV) Genome-browser, (V) Download, (VI) Statistics, (VII) Submit and (VIII) Help (**Figure 2A**).

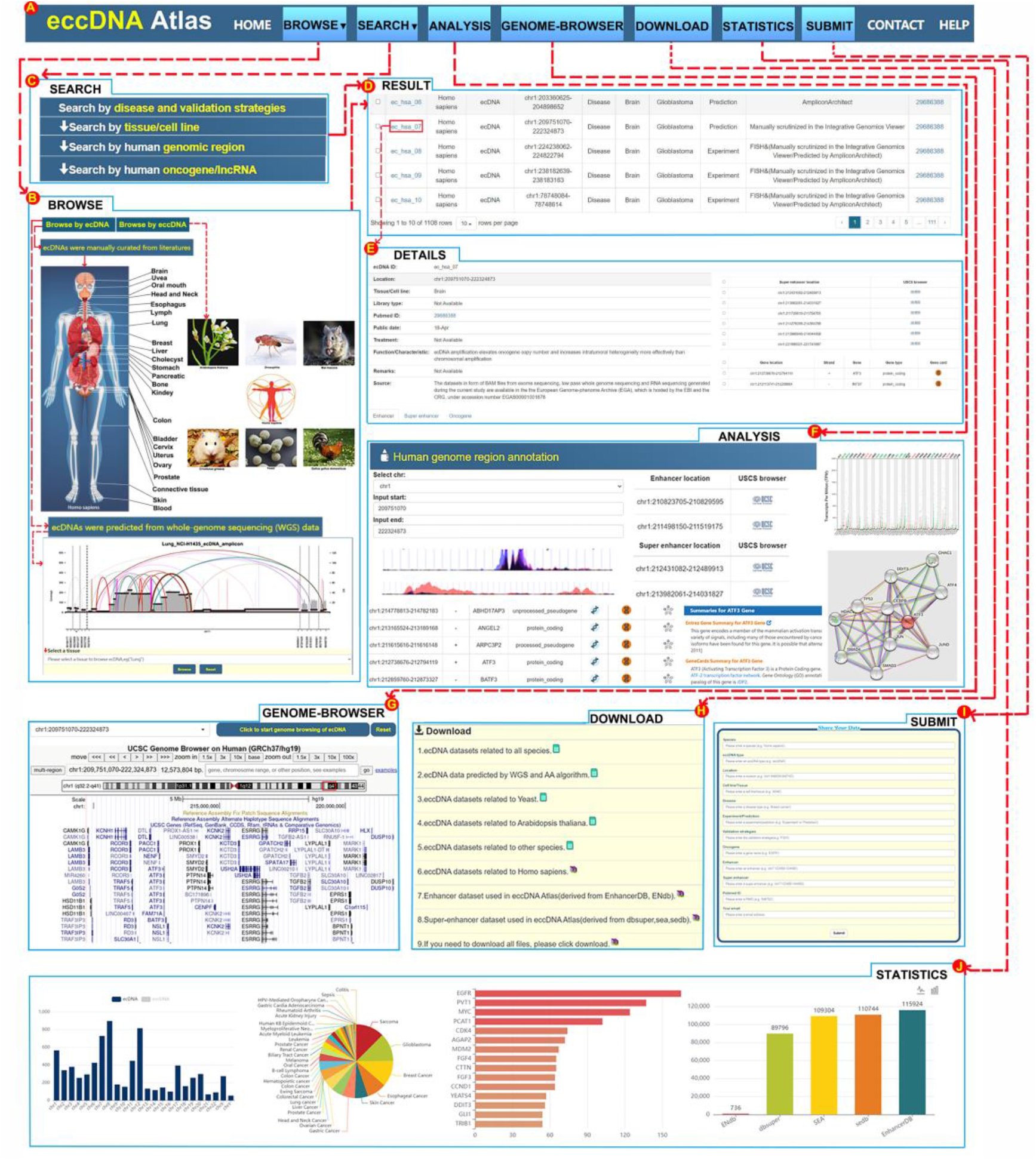

#### BROWSE

The browse page was divided into two menus: “browse by ecDNA” and “browse by eccDNA”. The “browse by ecDNA” page includes “manually curated from literatures” and “predicted from WGS data”. To make browsing easier for users, we classified the cell lines according to affiliated tissue source. Users can browse the ecDNAs by clicking the labels matching them up with the tissue type of schematic diagram or by choosing from the drop-down menu of tissues in the corresponding species catalog. In the eccDNA part, users can preferentially select species (eg: Drosophila, Yeast, Mus musculus, etc.) to browse eccDNA data, or choose the corresponding tissue to browse.

#### SEARCH

The search page was divided into two major parts: “search by ecDNA” and “search by eccDNA” (**Figure 2C**). Each part we further divided into four sections, including: 1) search by disease and validation strategies, 2) search by cell line/tissue, 3) search by genomic region, 4) search by oncogene/lncRNA. For each search section, we use a step-by-step guide for user-friendly searching. For example, for “search by disease and validation strategies”, the first step is to choose a species type. Secondly, a disease needs to be selected to refine the further search. Based on the search results of step 1 and step 2, experiment/prediction and validation strategies can be selected as an option (step3 and step4, respectively) for a more advanced search.

#### ANALYSIS AND GENOME-BROWSER

Users can perform annotation analysis on ecDNA region of eccDNA Atlas or custom chromatin regions including oncogenes/lncRNAs, typical enhancers and super enhancers. In addition, we provided some external links for gene expressions, functions, survival, and regulatory network exploring analysis, including GEPIA, GeneCard, and String databases (**Figure 2F**). Furthermore, we also provided genome visualization utilities for ecDNA based on UCSC Genome Browser. Users can select an ecDNA list of eccDNA Atlas or enter the user-defined ecDNA chromosomal coordinate range of ecDNA in the genome browse section, and get a general overview of the information contained in the ecDNA region such as histone modification signals, mutations, sequence conservatism, etc. (**Figure 2G**).

#### DOWNLOAD, SUBMIT AND STATISTICS

Users can download all eccDNAs/ecDNAs and annotation data from eccDNA Atlas (**Figure 2H**) or share their research data through submitting interface (**Figure 2I**). On the statistics page, we mainly count the oncogenes that occur frequently in eccDNAs/ecDNAs and the proportion or amounts of eccDNAs/ecDNAs in each disease or chromosome, etc. (**Figure 2J**).

#### A CASE STUDY

Take ec_hsa_6581 from prostate cancer as an example to introduce the use of eccDNA Atlas. In the browse page, users can click the prostate label to get the data containing ec_hsa_6581. In the search page, for example, for “search by disease and validation strategies”, users can first select the species type as “Homo sapiens” and secondly select the disease type as “Prostate Cancer” to get the data containing ec_hsa_6581. The result page contains the following information, including eccDNA ID, species, eccDNA type, location, health/disease, tissue/cell line, disease name, experiment/prediction, validation strategies, and pubmed ID. The details include publication date, treatment, function/characteristic, remarks and data source. Two frequently amplified oncogenes (MYC and PVT1) in human prostate cancer were annotated on ec_hsa_6581 by performing analysis of eccDNA Atlas, which are consistent with the previous study[13]. The results showed that PVT1 had a significant effect on patient survival in prostate cancer, and there was a relatively significant positive correlation between MYC and PVT1 (R=0.32, p=1.4e-12). In addition, ten typical enhancers and three super enhancers were annotated in this region, most of the enhancers had strong H3K27Ac signal peaks by genome-browser visualization of eccDNA Atlas. In summary, this information obtained from the database has important implications for the subsequent studies of the potential functions and regulatory ability of eccDNAs/ecDNAs.

## Discussion

Although EccDNA was discovered nearly 60 years ago, the eccDNA/ecDNA sequence information is insufficient in earlier research literatures due to technical limitations. In addition, studies that only reported the discovery of eccDNA instead of its function were excluded from our database. By the application of high-throughput DNA sequencing techniques, especially next-generation sequencing, it is possible to identify more and more eccDNAs. We collected as much as possible literatures containing eccDNA/ecDNA information at present by manually curated 3,636 literatures. As the most important class of eccDNAs, the mechanisms of ecDNA biogenesis are still unclear, and it is difficult to identify the high amplification region of ecDNA sequence with traditional experimental methods such as DNA electrophoresis, FISH, etc. In order to help researchers to study ecDNA more deeply, we collected massive WGS data from 319 cell lines by NCBI SRA and predicted 1,105 ecDNAs from 150 cell lines by AmpliconArchitect. Furthermore, we included a lot of annotation information for these ecDNAs such as oncogenes, lncRNAs, typical enhancers, super enhancers, etc. However, since there is no accurate method currently to distinguish whether the sequencing data come from linear chromosomes or eccDNAs, there are still many issues needed to be solved for accurate and comprehensive annotations of ecDNAs by current multi-omics data such as chromatin accessibility data, epigenetic data, Hi-C data, transcriptome data, etc.

In the future, we will continue to follow up on eccDNAs/ecDNAs related literatures. With the further amounts of data yielded by eccDNAs/ecDNAs, we will continuously collect the latest datasets to keep our database up-to-date. We will develop new database analysis functions, such as sequence alignment, and enhance the interactivity of the database to give users a better experience. In addition, with the advancement of experimental methods, it is expected to collect more sequencing data performed for eccDNAs instead of linear chromosomes, such as chromatin accessibility data on eccDNAs, 3D genome data between eccDNAs or eccDNA hubs, survival data, epigenetic modification data, etc. Overall, eccDNA Atlas is a comprehensive data portal of eccDNAs/ecDNAs for health and diseases across multiple species.

## Availability

The data are freely accessible for research purposes at http://lcbb.swjtu.edu.cn/eccDNAatlas/index.html

## Acknowledgments

We thank the Informatization and Network Management Office of Southwest Jiaotong University for providing the cloud-based technical support.

